# An Aggregation Prone Region (APR) in talin controls talin self-interactions to regulate integrin adhesion complex dynamics

**DOI:** 10.64898/2026.01.13.699108

**Authors:** Neil J. Ball, Paula Turkki, Yanyu Guo, Mahmudul H. Tanvir, Shimin Le, Jasmine Fei Li Chin, Karen B. Baker, Jingxing Luo, Nicholas H. Brown, Vesa P. Hytönen, Jie Yan, Benjamin T. Goult

## Abstract

The tight regulation of the integrin family of extracellular matrix (ECM) receptors is essential for the coordination of most cellular processes. Talin regulates integrin function by binding to the cytoplasmic tail of the integrin beta subunit. Two integrin-binding sites (IBS) in talin have been reported, one in the talin head domain (IBS1), and a second in the C-terminal rod region of talin (IBS2), mapped to the R11-R12 domains. Whilst the structural details of integrin binding to IBS1 are well understood, IBS2’s mode of binding integrin is less clear, and the biochemical details of this site have been elusive. Here we report that talin R11 contains a cryptic high affinity talin-binding site. We show that mechanical unfolding of R11 exposes an Aggregation Prone Region (APR) in Helix50 that oligomerizes with other talin R11s. Our data support a model whereby the process of mechanically unfolding one R11, exposes and maintains the APR in a high affinity conformation that drives unfolding of other R11 domains and oligomerisation. Atomic Force Microscopy confirms that the APR region alone forms micron long amyloid-like fibres, via a beta-sheet amyloid arrangement which can be abolished using a canonical “gatekeeper” point mutation, V2078K, that prevents APR interactions. Introducing this gatekeeper mutant in talin eliminates recruitment of talin IBS2 to integrin adhesion sites. The APR region in talin overlaps with a vinculin-binding site in Helix50, and we show a novel role for vinculin as a mechano-chaperone for talin, able to resolve the talin large species by binding to the VBS in Helix50. In light of this new information, we propose that the ability of R11-R12 to target to integrin adhesion complexes and enhance integrin activation can be better explained by a model where functional aggregation of talin serves to recruit and activate more talin molecules at the adhesion site leading to enhanced integrin activation. Whilst further work is required to fully exclude integrin binding to R11 we suggest that the name IBS2 might be a misnomer and propose the name Aggregation Prone Region 1 (APR1) for this talin oligomerisation motif.

## Introduction

The adhesive structures that cells make to attach to the extracellular matrix (ECM) are beautiful in their sophistication and complexity, able to form numerous different signalling complexes and structures as a function of the properties of the ECM. The integrin family of ECM-receptors mediate this adhesion and large integrin adhesion complexes (IACs) assemble at these sites, coupling the ECM to the cytoskeleton via the adaptor protein talin. A significant advance has been the realisation that, despite their structural complexity (containing potentially hundreds of proteins (Horton et al., 2016)), these adhesion complexes contain a simple and robust core of just four proteins: integrin, talin, kindlin and actin (Goult et al., 2021; Theodosiou et al., 2016).

Talin is a large multidomain protein, that comprises an N-terminal FERM domain (with subdomains F0-F3), followed by an ∼80 aa unstructured linker, and then 13 helical bundles, R1-R13 and a dimerisation helix (DD), which form the talin rod (Gingras et al., 2008; Goult et al., 2013). Each rod domain has the potential to function as a mechanochemical switch, as it can reversibly switch between folded “0” and unfolded “1” states in response to mechanical forces (Fig.1A) (Goult et al., 2013; Haining et al., 2016; Yao et al., 2016, 2014). These different states can recruit different proteins to alter IAC composition (Ball et al., 2024; Goult, 2021). Many talin-interacting proteins have been identified (Goult et al., 2021), emphasising the ability of talin to function as a mechanosensitive signalling hub (Goult et al., 2018).

**Figure 1.**
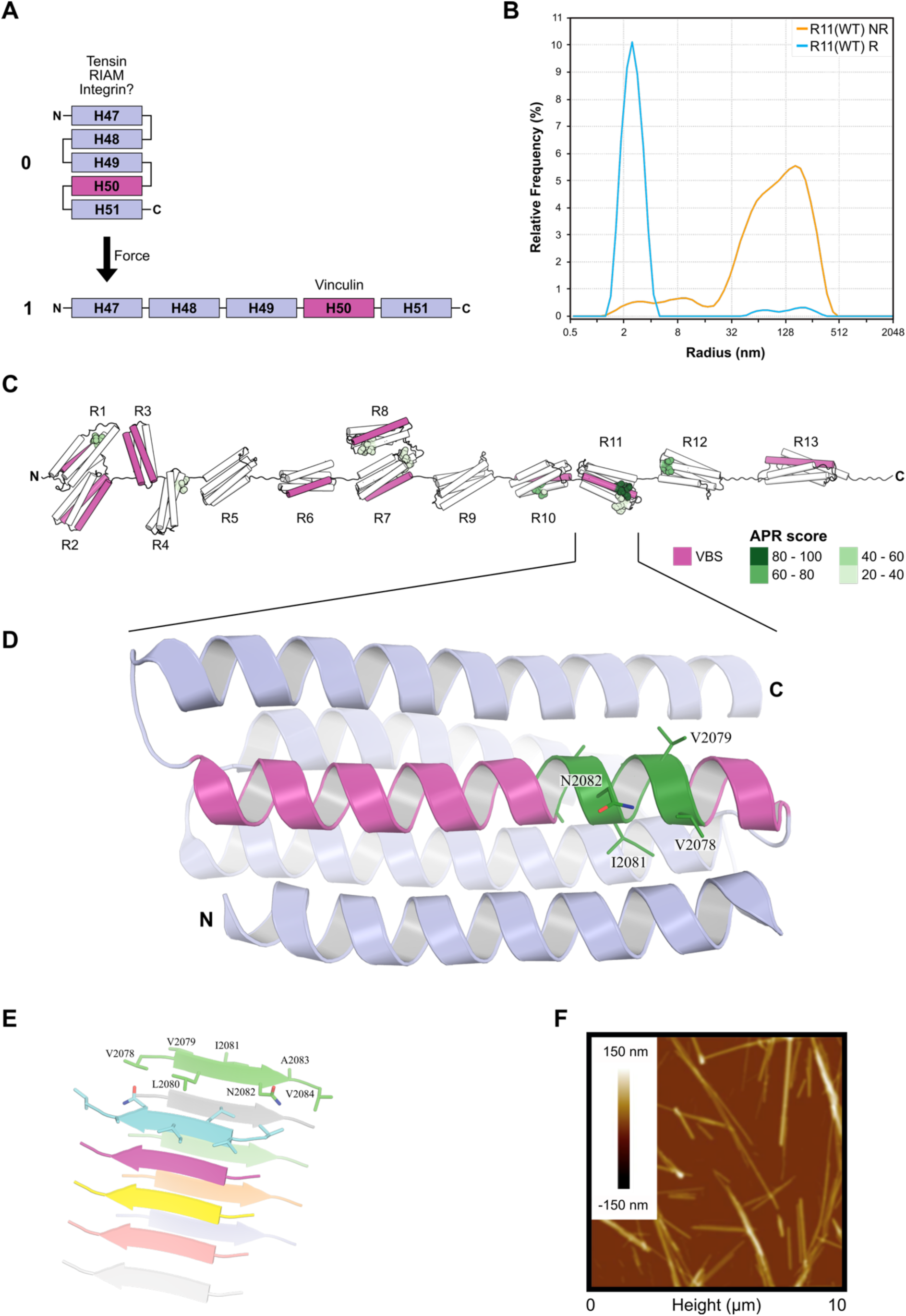
R11 contains an Aggregation Prone Region (APR) that can form amyloid-like fibrils *in vitro*. **(A)** Cartoon of the R11 five-helix bundle of talin in the folded “0” and unfolded “1” states with the known interactors labelled. Helix50 contains a vinculin-binding site (VBS) shown in magenta. **(B)** Talin R11 can form a big species. Dynamic Light Scattering (DLS) of reduced (blue) and non-reduced (orange) R11. The non-reduced sample of R11 has a large hydrodynamic radius (∼128 nm) indicating the presence of a large soluble multimer. Addition of a reducing agent resulted in loss of the larger species reverting to a single peak at 2.47 nm which corresponds to monomeric R11. **(C)** Location of predicted APRs in talin1. The predicted APRs are shown as spheres and coloured according to their APR score (green) as calculated by the TANGO algorithm (A.-M. Fernandez-Escamilla et al., 2004). The highest scoring APR is in R11 (dark green). Vinculin binding sites (VBSs) are shown in magenta. **(D)** The APR in R11 (green), sequence ^2078^VVLINAV^2084^, overlaps with the VBS in Helix50 (magenta). **(E)** AlphaFold structural model of 10 copies of the VVLINAV APR region from R11. The APR is predicted to form beta strands that can self-assemble into beta sheets, which are then stacked. **(F)** Atomic Force Microscopy (AFM) image of a ^2078^VVLINAV^2084^ synthetic peptide that has self-assembled *in vitro* to form extended micron-long amyloid-like fibrils.

Forming a mechanical linkage between the ECM-integrin complex and actin involves direct interactions between talin, integrin and actin. The binding of talin to integrin can potentially be mediated by either of two reported integrin-binding sites (IBSs) (Calderwood et al., 2013; Calderwood, 2004; Critchley, 2009). The direct binding of IBS1 to the cytoplasmic tail of the integrin beta subunit has been well established by numerous experimental approaches (reviewed in (Calderwood, 2004)), including atomic structures of the complex (Anthis et al., 2009), however for IBS2, the evidence is less equivocal. The presence of an integrin binding site in the rod domain was first described as integrin binding to the two talin fragments formed by calpain cleavage (head and rod) (Horwitz et al., 1986). Although the data for the rod binding integrin was not shown it suggested that there were at least two integrin-binding sites in talin. This was followed by demonstration that a region of the talin rod (residues 1984-2541; comprising part of R11, R12, R13) bound to integrin β3 coated wells (Xing et al., 2001). A region of the talin rod that was sufficient for localisation of rod fragments to focal adhesions (FAs) was mapped to a 130-residue region, and then refined to a 23 residue fragment (residues 2077-2099), corresponding to helix 50 of R11 (Moes et al., 2007). The structure of R11-R12 showed two 5-helix bundles sharing a long alpha helix, and both R11-R12 were found to be required for integrin binding (Gingras et al., 2009). A common feature of the biochemical assays used to demonstrate IBS2 binding to integrin is that they all used a solid phase approach (Gingras et al., 2009; Moes et al., 2007; Xing et al., 2001), with the integrin cytoplasmic domain attached to a surface. Binding of IBS2 to talin in solution has yet to be reported.

An interaction between IBS2 and integrin has also been supported by genetic experiments in *Drosophila*. Point mutations in either IBS1 or IBS2 impair talin function, and the double mutant is stronger, indicating some compensatory function between the two IBSs (Ellis et al., 2011). Analysis of a large number of truncation mutants of talin, revealed that Helix50 was particularly important in mediating adhesion between epithelial layers, but not for mediating muscle attachment to tendon matrix (Klapholz et al., 2015). Using FRET, integrin and IBS2 were found in close proximity in the epithelium, but not the muscle, and the IBS2s in different talin molecules were found close together *in vivo*.

In this study, we report the discovery of an aggregation prone region (APR) in talin, which maps to Helix50 in R11. We show that the oligomeric status of R11 is very conformationally sensitive, and changes in force and/or redox conditions can expose this APR and trigger oligomerisation. We show that Helix50 is involved in three different interactions, i) in the folded “0” state the R11 bundle is involved in binding to the ligands tensin-3 (Atherton et al., 2022) and RIAM (Goult et al., 2013), ii) in the unfolded “1” state where the domain is unfolded to a string of helices, Helix50 binds vinculin (Gingras et al., 2009, 2005; Yogesha et al., 2012), and iii) when unfolded to an unstructured polypeptide it forms a high affinity binding site for other Helix50s. We propose the name Aggregation Prone Region 1 (APR1) for this talin oligomerisation motif. In this way we suggest that there may not be a second integrin-binding site in the talin rod and instead the data supporting IBS2 binding to integrin can be re-interpreted as talin oligomerisation. This leads to a new model, whereby unfolding of R11 mediates functional aggregation of talin, clustering talin molecules and stabilising their active conformation, which in turn leads to increased integrin activation. Finally, we show that vinculin binding can resolve these R11 oligomers, thus revealing a novel role for vinculin as a mechanochaperone.

## Results

### Talin R11 can form higher order species

Expression and purification of recombinant talin R11 (residues 1974-2140) gives rise to a well behaved soluble protein (Gingras et al., 2009). However, we noticed that over time the protein can undergo changes in its properties. This can be visualised in Fig.S1A where partially purified R11 protein was processed in two different ways prior to loading onto an anion exchange column. When immediately diluted with 20 mM Tris pH 8.0, 50 mM NaCl, a single peak is observed (light blue in Fig.S1A), whereas after dialysis overnight into the same buffer the elution profile contains two peaks (dark blue in Fig.S1A). Analysis of these two peaks by reducing SDS-PAGE revealed both peaks contain the same protein (Fig.S1B). However, the use of non-reducing sample buffer revealed that the second peak is twice the size of the monomeric protein, suggesting that R11 forms a disulphide-bonded dimer (Fig.S1B). R11 contains a single cysteine residue, Cys1978 that is at the start of the first helix (Helix47) and is buried in the core of the folded R11 domain (Fig.S1C). The broad nature of the additional non-reduced ion exchange peak led us to examine it by Dynamic Light Scattering (DLS) to look at its hydrodynamic radius (Fig.1B). DLS revealed that the non-reduced R11 sample has a very large hydrodynamic radius (∼128 nm), which is dramatically larger than a dimer (Fig.1B, orange). Addition of the reducing agents Dithiothreitol (DTT; not shown), or Tris(2-carboxyethyl)phosphine (TCEP; Fig.1B, cyan) fully converted the large species back to the monomeric form, as analysed by DLS and SDS-PAGE. The estimated molecular weight of a protein with hydrodynamic radius of 128 nm is 800 MDa, corresponding to a multimer with approximately 40,000 copies. This indicates that the single buried cysteine residue in R11 forms a disulphide bond between two R11 monomers, and stabilises a form of R11 that is able to form large soluble aggregates of R11. This can be explained if forming the disulphide bond requires partial unfolding of the domain, and the disulphide bond stabilises this unfolded state, which then forms large aggregates. We note that the term “aggregation” is often incorrectly thought of as solely a negative behaviour of ill behaving proteins, but functional protein aggregation is a recognised phenomenon in many systems (Gsponer and Babu, 2012).

### Computational analysis identifies that talin contains multiple aggregation prone regions (APRs)

The large species of R11 that can form once R11 is partially unfolded indicated that the domain contains a cryptic talin interaction motif. To look for this motif we used the TANGO software (A. M. Fernandez-Escamilla et al., 2004), which predicts Aggregation Prone Regions (APRs) in proteins. These APRs have propensity to form intermolecular beta-sheets and are established forms of protein oligomerisation, best known for their role in pathogenic amyloid formation (Riek, 2017), but many examples of functional APR motifs have been reported. TANGO analysis of talin1 revealed multiple APR regions distributed through the talin rod (Fig.1C) and it was striking that these APRs are predominantly located buried within the folded talin rod domains (Fig.1C). In particular, the highest scoring APR in talin, corresponds to the sequence ^2078^VVLINAV^2084^ in R11. This sequence is part of the 4^th^ helix of the 5-helix bundle and forms part of the vinculin-binding site (VBS) in Helix50 (Fig.1D). The crystal structures of R11-R12 (pdb 3dyj (Gingras et al., 2009)) and Helix50 in complex with vinculin D1 domain (VD1) (pdb 4dj9 (Yogesha et al., 2012)) are both solved and show this APR region is an integral part of the helix in both the folded talin R11 bundle and when in complex with VD1.

### AlphaFold structural modelling indicates that the APR region of R11 can form intermolecular beta-sheet structures

To further explore the potential of this VVLINAV region to form oligomers we used AlphaFold3 (Abramson et al., 2024) to generate a structural model of 10 copies of a VVLINAV peptide (Fig.1E). The resultant structural model predicted the beta-sheet amyloid arrangement of the 10 talin peptides, with each VVLINAV motif forming a beta-strand that stacks into a parallel beta-sheet with these beta-sheets stacked together in an anti-parallel orientation.

### The isolated APR from R11 forms micron long amyloid fibres, which can be eliminated by a V2078K gatekeeper mutation

To validate this predicted APR region, we synthesised two peptides, ^2078^VVLINAV^2084^, and a version where we introduced a “gatekeeper” residue, mutating V2078 to lysine, ^2078^**K**VLINAV^2084^. Gatekeeper residues are charged residues that are positioned at the start of an APR such that stacking of repeating peptides is disfavoured due to the charge repulsion (Beerten et al., 2012). We chose the first valine, Val2078, to mutate due to it being solvent exposed in the structure of R11 (Fig.1D). The VVLINAV peptide readily formed fibrous structures which were visible to the naked eye. When visualised with Atomic Force Microscopy (AFM), often used to examine amyloid structures (Adamcik and Mezzenga, 2012), these fibres were observed to be microns long and of consistent diameter (Fig.1F). As predicted, the gatekeeper V2078K version eliminated the formation of fibres (no data to show).

### The cysteine is not required for R11 self-assembly if R11 is destabilised

Both cysteine 1978, and the APR region in R11 are cryptic, buried in the folded R11 domain (Fig.S1C and 1D respectively) and so are inaccessible to interactions without a conformational change. This suggested that factors that induce conformational changes in R11 may trigger oligomerisation. The formation of the disulphide-mediated R11 dimer appears to be necessary for the formation of larger species of R11 *in vitro*, as these do not form in a reducing environment. This led us to test three hypotheses, i) the cysteine is required for big species formation, ii) R11 dimerisation is required for big species formation and iii) it is the conformational change that is required for big species formation. To test these hypotheses, we made a series of new constructs of R11. First, we mutated the cysteine in R11 to alanine (C1978A). The R11(C1978A) protein behaved well and produced a monodisperse single species in DLS in both reducing and non-reducing conditions (Fig.S1D; orange) with a hydrodynamic radius equal to that of a monomer, identical to the reduced R11(WT) (Fig.1B; light blue). This confirmed that *in vitro* the cysteine is necessary for large species formation of R11. Next, we wanted to test whether disulphide-mediated dimerisation by itself is sufficient for aggregate formation so we designed a further construct where we added a cysteine attached via a short linker to the C-terminus of R11(C1978A) (R11(C1978A)-Cys, Fig.S1F; inset). R11(C1978A)-Cys readily formed dimers in non-reducing conditions (Fig.S2B) but was unable to form large species, supporting the idea that it is not dimerisation of the R11 itself but the disulphide-mediated stabilisation of the unfolded species of R11 that is required. Finally, to test the requirement of conformational change to expose the APR motif, we made an R11 construct with a deletion of the cysteine-containing first helix (Helix47; residues 1974-2009). Deleting the first helix both removes the cysteine and destabilises the domain. This truncated, R11(ΔH47) protein readily formed large aggregates, even in reducing conditions (Fig.S1F; dark blue). Taken together, this data points to R11 unfolding as the important trigger for self-assembly, a process accelerated by the disulphide-mediated stabilisation of the unfolded conformation.

### Talin R11 is only recruited to cell adhesions formed by R11-containing talin

Having established that self-interactions of R11 occur *in vitro,* we next wanted to explore the role of R11-R11 interactions at focal adhesions in cells. Integrin adhesion complexes require the presence of talin to mediate the mechanical linkage between the integrin-ECM complex and the cytoskeleton. Removal of talin results in cells rounding up and detaching (Theodosiou et al., 2016) as exemplified in *Tln1⁻/⁻Tln2⁻/⁻* MKF cells that are round and non-adherent. Transfection of these cells with wild type talin constructs restores their ability to adhere and form focal adhesions (FAs) (Fig.2A). This makes them ideal for looking at specific talin functions as they lack endogenous talin, which would otherwise complicate the analysis of modified talin constructs.

**Figure 2.**
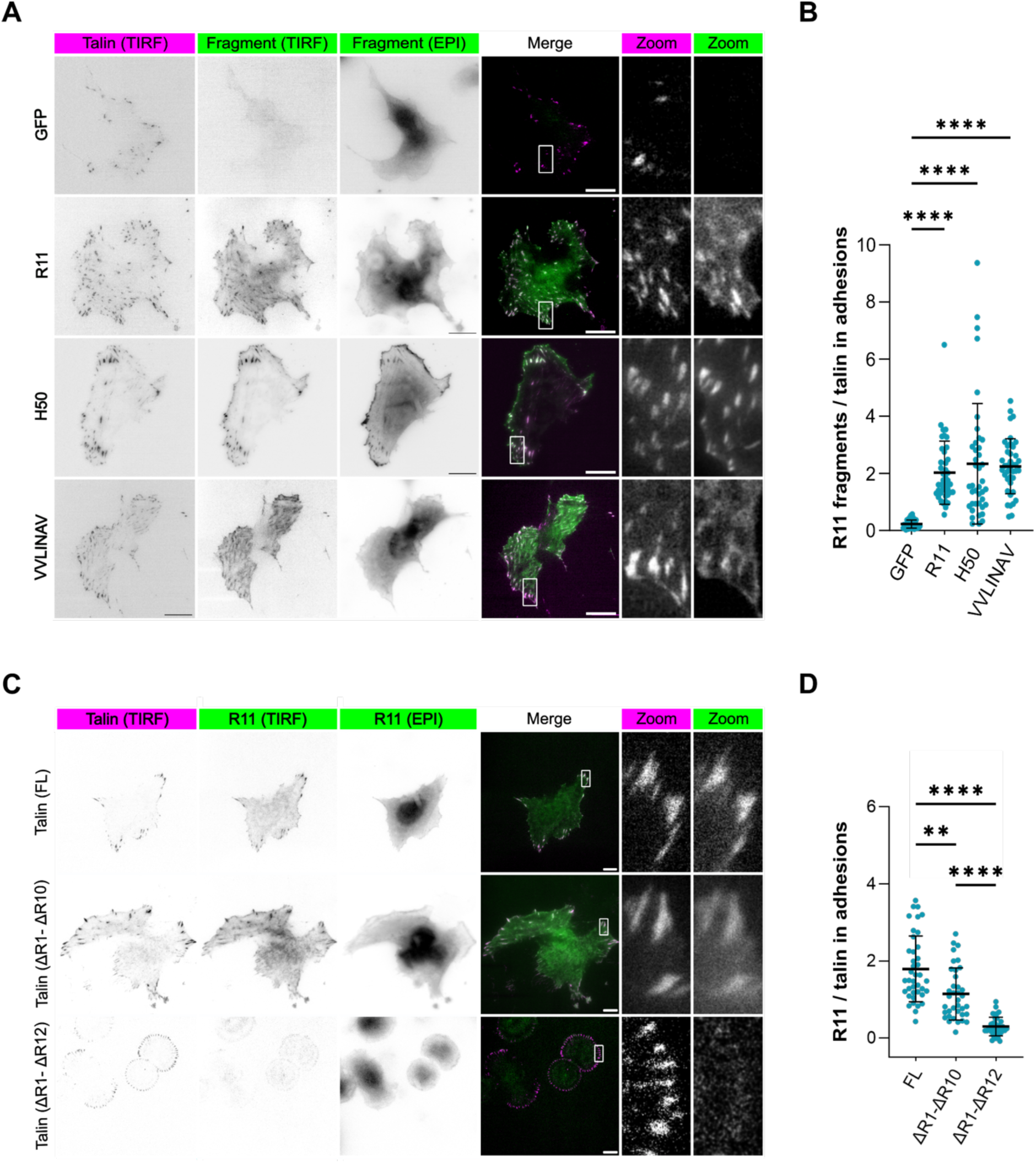
Recruitment of talin R11 to cell adhesions requires talin with R11 to be already present. **(A)** Representative TIRF images showing colocalisation of GFP-R11 fragments (GFP-R11, GFP-H50 and GFP-VVLINAV) with mCherry full-length talin1 (Talin(FL)). Scale bar 10 µm. **(B)** Quantitative analysis of R11 fragment colocalisation at adhesion sites with Talin(FL) (two independent experiments n =37-41 cells). Colocalisation within adhesions was quantified as the GFP intensity ratio relative to mCherry intensity. **(C)** Representative TIRF images showing colocalisation of the R11 domain with full-length talin and talin lacking domains R1–R10, Talin(ι1R1-R10) but not with talin lacking domains R1-R12, Talin(ΔR1–R12). Scale bar 10 µm. **(D)** Quantification of R11 domain colocalisation at adhesion sites for the indicated talin constructs (three independent experiments n =30-40 cells). A similar analysis for GFP-VVLINAV is shown in Fig.S2. Statistical significance was assessed using Brown–Forsythe and Welch ANOVA tests. Data are presented as mean and SD from three independent biological replicates. Statistical significance is indicated as follows: *P < 0.05; **P < 0.01; ****P < 0.001.

We generated three EGFP-tagged R11 constructs, namely GFP-R11, GFP-Helix50 (GFP-H50) and GFP-VVLINAV and co-expressed them, or GFP alone as a control, with mCherry/mScarlet-labelled full-length talin (Talin(FL)). Upon expression of Talin(FL) the cells spread and made FAs. Co-expression of GFP-R11, GFP-Helix50 and GFP-VVLINAV revealed that all three R11 constructs localised to the mCherry-Talin(FL) containing adhesions (Fig.2A, quantified in Fig.2B).

The localisation of GFP-R11 and GFP-H50 to FAs was in good agreement with previous work (Moes et al., 2007; Tremuth et al., 2004) which mapped the FA-targeting region in R11 to residues 2077-2099. However, as the GFP-VVLINAV sequence is sufficient to target to FAs we have further refined this FA-targeting motif in talin to just seven residues (2078-2084). In summary, we show that the APR region in talin is sufficient to localise to FAs.

Having established that GFP-R11 fragments can target to FAs containing Talin(FL) we next wanted to test whether deletion of R11 from Talin(FL) would reduce GFP-R11 targeting, using mini-talin constructs lacking most of the talin rod: Talin(ýR1-R12) and Talin(ýR1-R10) (Rahikainen et al., 2019). Upon transfection, these mini-talins enable cell spreading of the talin null cells. GFP-R11 co-localised with Talin(FL) and Talin(ýR1-R10) but did not co-localise with Talin(ýR1-R12) which lacks R11-R12 (Fig.2C,D). The absence of co-localisation was also seen for GFP-H50 (data not shown) and GFP-VVLINAV (Fig.S2A,B). In summary, we show that the APR region in talin R11 is required to localise R11 to FAs.

### The canonical APR-disrupting gatekeeper mutation, V2078K, prevents R11 self-assembly *in vitro* and FA-targeting of R11 in cells

The APR-motif in talin R11 by itself forms micron long fibres (Fig.1F), and this self-association can be abolished by introduction of a gatekeeper point mutant, V2078K. Having validated the effectiveness of the V2078K mutation in limiting talin amyloid formation at the peptide level, we next introduced this mutation into the R11 domain. R11(V2078K) expressed and purified well, and DLS analysis confirmed that R11(V2078K) no longer forms the large species (Fig.3A). This demonstrates that the V2078K mutation is effective as a gatekeeper and is able to limit talin aggregation in the context of the R11 domain.

**Figure 3.**
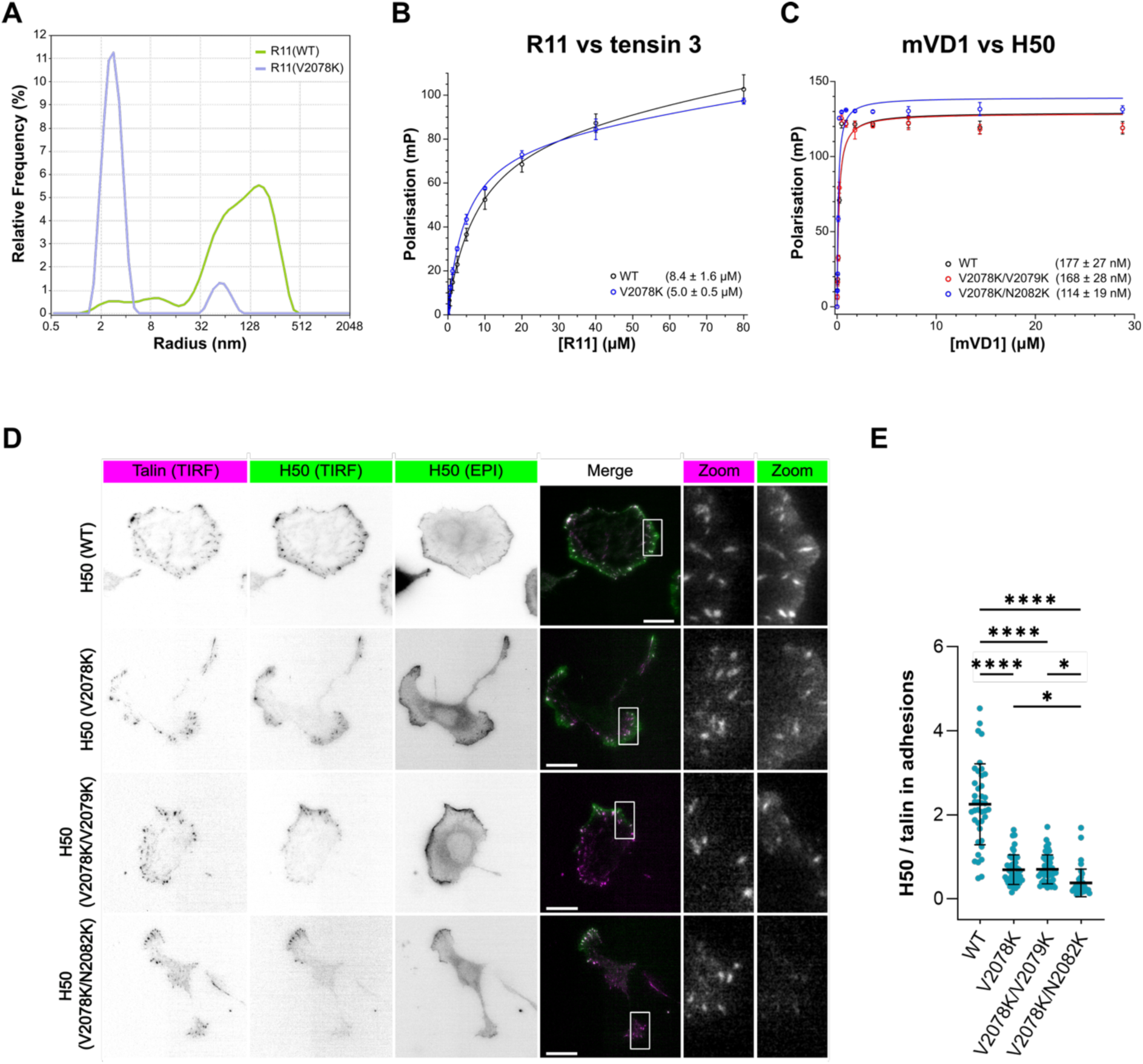
A gatekeeper mutation, V2078K, in the APR prevents R11 self-assembly *in vitro* and in cells. **(A)** DLS comparison of R11(WT) (green) and R11(V2078K) (blue) in the absence of reducing agent. The gatekeeper mutation significantly reduces the propensity for R11 to self-assemble. **(B-C)** The V2078K gatekeeper mutation does not affect other R11 interactions. **(B)** Fluorescence polarisation of WT (black) and V2078K (blue) binding to a tensin-3 peptide (K_D_ shown in parentheses). All proteins were fully reduced with 0.5 mM TCEP prior to measurements. **(C)** The V2078K (gatekeeper) mutation is in the VBS in Helix50 but does not affect binding to vinculin VD1 domain. **(D)** Representative TIRF images showing colocalisation of wild-type Helix50 and Helix50 gatekeeper mutants (V2078K, V2078K/V2079K, V2078K/N2082K) with full-length talin. Scale bar 20 µm. **(F)** Quantitative analysis of colocalisation between wild-type Helix50 or gatekeeper mutants and full-length talin (three independent experiments n =35-40 cells). Statistical significance was assessed using Brown–Forsythe and Welch ANOVA tests. Data are presented as mean and SD from three independent biological replicates. Statistical significance is indicated as follows: *P < 0.05; **P < 0.01; ****P < 0.001.

R11 interacts with multiple proteins (Fig.1A), and contains binding sites for tensin (Atherton et al., 2022) and RIAM (Goult et al., 2013) when folded, and vinculin when unfolded (Gingras et al., 2005; Yogesha et al., 2012). Therefore, before using this mutation for cellular analysis, we first confirmed that the V2078K mutation did not affect binding to tensin-3 (Fig.3B) or VD1 (Fig.3C).

Having validated the efficacy of the V2078K mutant in biochemical assays, we introduced the gatekeeper mutation to our GFP-H50 construct, GFP-H50(V2078K), and studied the effect on colocalisation with full-length talin at cell adhesions (Fig.3D). The mutant dramatically reduced the colocalisation of GFP-H50 to focal adhesions (Fig.3E), although some FA-localisation was still observed. Therefore, we tested two further mutants, V2078K/V2079K and V2078K/N2082K, that were designed to further limit self-assembly by further enhancing the charge repulsion of the V2078K. Both mutations still bound to vinculin *in vitro* (Fig.3C) and both further reduced the colocalisation of GFP-H50 with talin at cell adhesions (Fig.3D,E) with the V2078K/N2082K being the most effective. This analysis validated the efficacy of the V2078K mutation at markedly reducing self-interactions *in vitro* and in cells.

### Changes in cell morphology, migration and adhesion component recruitment due to V2078K mutation in full-length talin

Having validated the V2078K mutant as an effective way to reduce talin interactions, we next wanted to understand the biological relevance of these talin self-interactions in cells. To do this we introduced the V2078K gatekeeper mutation into full-length talin and evaluated the effects on cell morphology and adhesion composition relative to the wildtype. We transfected *Tln1⁻/⁻ Tln2⁻/⁻* MKF cells with wild type talin or the V2078K mutant. Cells expressing the V2078K mutant were larger (area 1539 ± 686.5 μm^2^) as compared to those expressing wild type talin (area 1315 ± 639.6 μm^2^) while aspect ratio (V2078K, 1.81, ± 0.53; WT, 1.76, ± 0.61) and circularity (V2078K, 0.28, ± 0.17; WT, 0.29, ± 0.18) showed no difference (Fig.4A,B). Additionally, the V2078K expressing cells showed a more centralised adhesion pattern, compared to wild-type cells where adhesions were more prominently localised at the cell edge (Fig.4C).

**Figure 4.**
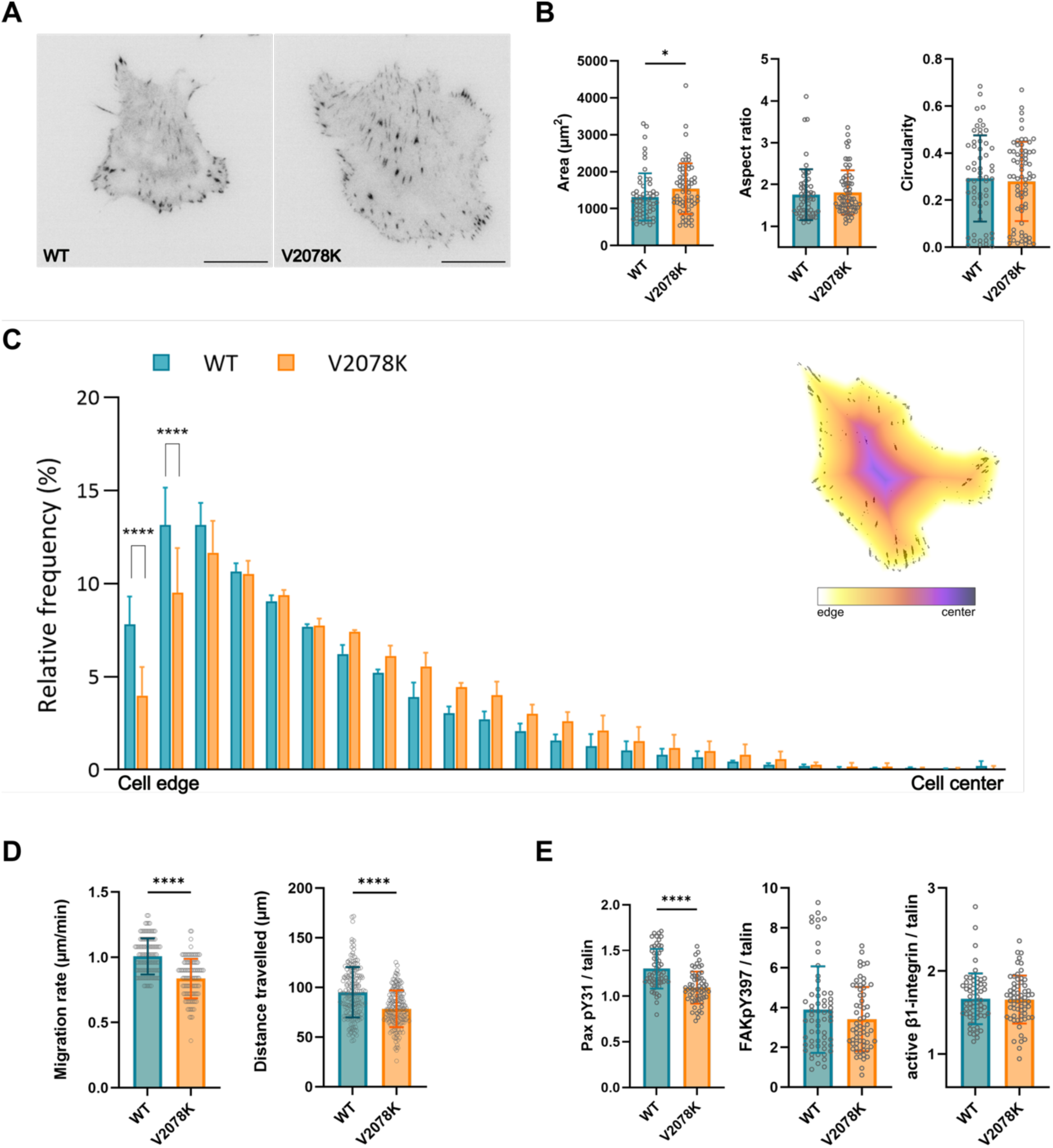
Changes in cell morphology, migration and adhesion component recruitment due to the V2078K mutation in full-length talin. **(A-B)** Quantitative analysis of cell morphology parameters including cell area, aspect ratio, and circularity in V2078K talin-transfected cells compared to wild-type (WT) talin (three independent experiments n=76-67 cells for each). Statistical significance was assessed using the Mann–Whitney U test. **(C)** Quantification of adhesion localisation in respect to cell edge and cell centre. Statistical significance was evaluated using one-way ANOVA. Data are presented as mean and SD from three independent biological replicates. **(D)** Random migration analysis including migration rate (μm/min) (three independent experiments n=160-179 cells) and total distance travelled (μm) (three independent experiments n=175 cells) for V2078K talin-and wild-type (WT) talin transfected cells. Statistical significance was determined by Welch’s t-test. (E) Quantification of adhesion-associated signalling molecules, represented as intensity ratios of paxillin pY31, FAK pY397, and activated β1-integrin at adhesion sites (three independent experiments n=60 cells for each). Mann–Whitney U test was used for statistical evaluation. Statistical significance is indicated as follows: *P < 0.05; **P < 0.01; ****P < 0.001.

V2078K-expressing cells also showed reduced migration rate (0.84 ± 0.15 μm/min) and slower wound closure (78.43 ± 18.44 μm) relative to wild-type (1.01 ± 0.14 μm/min; 95.06 ± 25.33 μm, respectively) (Fig.4D). To assess adhesion dynamics, we quantified known adhesion molecules at cell adhesions whose activation status can be quantified using antibodies (pY31 paxillin, pY397 FAK, and activated β1-integrin). V2078K-expressing cells showed significantly lower (1.09 ± 0.18) pY31 paxillin recruitment compared to wild-type (1.30 ± 0.22) whereas no differences were observed for pY397 FAK levels (V2078K, 3.42 ± 1.61; WT, 3.90 ± 2.17) or activated β1-integrin (V2078K, 1.66 ± 0.29; WT, 1.67 ± 0.31) (Fig.4E). In summary, the addition of a gatekeeper mutant into APR1 results in alterations in adhesion dynamics and reduced migration and wound closure consistent with perturbed talin function.

### Talin R11 oligomerisation is mechanically-gated

The finding that destabilising R11 is sufficient to trigger self-association (Fig.S1F) suggested that mechanical unfolding of the R11 domain could be a normal mechanism to mechanically regulate talin oligomerisation via the APR. Our previous work on other cryptic binding sites in talin, including the multiple vinculin binding sites that are exposed upon rod domain unfolding (Yao et al., 2016, 2014) and the A-kinase Anchoring Protein (AKAP) helix in R9 that binds Protein Kinase A when R9 is unfolded (Kang et al., 2024), revealed a way to assay binding to an unfolded domain at the single molecule level. When talin is put through cycles of stretch to progressively unfold each of the domains and relaxation to allow them to refold, addition of a ligand, e.g. vinculin, causes the loss of specific unfolding step(s), because the domain(s) bound by ligand in its unfolded state are unable to refold during the relaxation step.

To study the talin R11-talin R11 interaction we applied single molecule magnetic tweezer approaches to mechanically stretch talin R9-R12 as described previously (Kang et al., 2024; Yao et al., 2016) and used free talin R9-R12, R11 or H50 as the ligand (Fig.5A,B and Fig.S3). In this experiment, multiple force cycles are applied to the protein. Initially, a linearly increasing force (1.5-30 pN at a constant loading rate of 2 pN/s) is applied to the protein tether which unfolds the domains, before the force is lowered back to ∼1.5 pN to allow refolding. When stretching R9-R12 alone four unfolding steps are observed in each cycle (red arrows in Fig.5b), confirming that in the absence of other proteins, these domains exhibit switch like behaviour and unfold and refold with high fidelity. We next applied the same magnetic tweezer technique but added in R9-R12, R11 or H50 as a ligand into the experiment and watched how they effected the tethered R9-R12 behaviour (Fig.5B). Strikingly, addition of R9-R12 into the chamber resulted in the loss of one unfolded step (Fig.S3), confirming that one domain was unable to refold, indicating an interaction with the tethered R9-R12 protein (Fig.S3). A similar loss of unfolding step was seen when just R11 (Fig.5b) or H50 alone (Fig.S3) were added enabling us to ascribe the lost unfolding step to R11.

**Figure 5.**
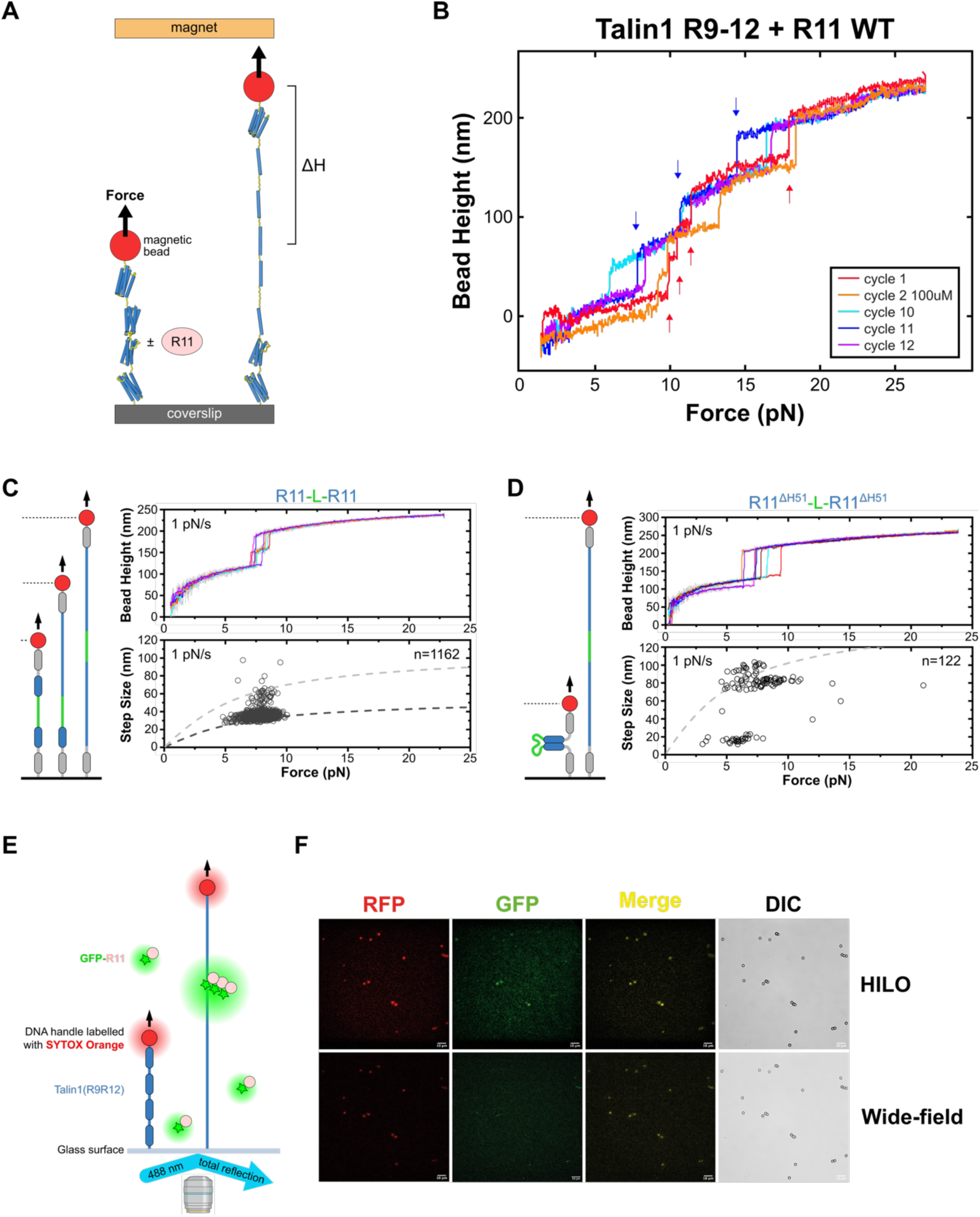
Mechanical gating of the APR in talin R11. **(A)** Schematic showing the single-molecule magnetic tweezers experiment. Talin R9-R12 is tethered between coverslip and a superparamagnetic bead using a 572–base pair DNA handle. Force is exerted and domain unfolding results in stepwise jumps in bead height (ΔH). **(B)** Representative consecutive force-height curves obtained from a tether of R9-R12 at a loading rate of 0.4 ± 0.04 pN/s are shown. The first cycle is R9-R12 alone and four unfolding steps are detected (red arrows). On cycle 2 100 µM R11 is added and within a few cycles one unfolding step is lost (blue arrows), once lost this step is lost in all subsequent cycles. **(C-D)** Schematics showing the single-molecule detector assay experiment. Two copies of R11 (blue; either R11 **(C)** or R11ΔH51 **(D)**) are connected via an unstructured polypeptide linker (green). Top: Force-bead height and Bottom: Force-step size curves for **(C)** R11-R11 and **(D)** R11ΔH51-R11ΔH51. The R11 domains refold faithfully each time and give two discrete unfolding steps. In contrast the R11Δ51 unfolds as a single unfolding step indicating a direct interaction between the two copies. **(E)** Schematic showing the single-molecule magnetic tweezers-TIRF experiment. Magnetic tweezers are installed over a TIRF microscope, and then the DNA handle is labelled with SYTOX Orange and can be visualised in the red channel. GFP-R11 and GFP-H50 are then added into the chamber. **(F)** Visualisation of multiple beads. Upon force addition, the green GFP signal grows at the same place as the red channel signal from the SYTOX Orange indicating coalescence of many GFP-R11 molecules onto the stretched R9-R12 seed.

With the single-molecule tweezer experiment we are detecting changes in the single molecule that is tethered, and so we cannot directly observe the ligand. With talin oligomerising via the APR we next wanted to test whether; i) two molecules of R11 are sufficient to interact, and ii) one mechanically-stretched R11 can recruit multiple R11 molecules from the surrounding solution. To test these scenarios, we used two variations of the single molecule experiments we had established in our lab.

### Two molecules of R11 are sufficient to interact

Measuring binding kinetics at the single molecule level is challenging as when the ligand dissociates it diffuses away making multiple repeats challenging. Therefore, we previously developed a single-molecule detector assay (Le et al., 2019) where the two proteins are connected via an unstructured polypeptide ensuring they faithfully rebind once the force is removed and refolding occurs. We made two versions of the experiment, one with two copies of R11 and the second with two copies of R11 lacking the last helix (R11ΔH51) to destabilise R11 (Fig.5C-D). The R11-R11 assay showed highly robust force-dependent unfolding and refolding of R11: the two R11 domains were able to unfold at around 7 pN and refold independently over many cycles with high fidelity (>95%) (Fig.5C). In contrast the R11ΔH51- R11ΔH51 assay showed only a single large rupture step (∼95 nm), or a small partial unfolding step (∼15 nm), followed by a large rupture step (∼80 nm), consistent with extension change of releasing the long linker upon unfolding. The results indicate that the two copies of the destabilised R11 interact consistently each cycle and rupture co-operatively (Fig.5D).

### One mechanically-stretched R11 molecule will recruit multiple R11 molecules from the surrounding solution

The final single molecule experiment we performed was to combine two mature technologies, namely single-molecule magnetic tweezers and total internal reflection fluorescence (TIRF) microscopy (Fig.5E). We also used highly inclined and laminated optical (HILO). TIRF excites molecules within 100-200 nm of the coverslip whereas HILO excites within a few microns. In this assay the R9-R12 tether was attached to a magnetic bead via a DNA handle that was fluorescently labelled with SYPRO orange and then stretched in the absence or presence of GFP-R11 or GFP-Helix50. The tether is then observed using TIRF/HILO and Differential Interference Contrast (DIC) microscopy to visualise the bead, the tethered SYPRO Orange labelled R9-R12 molecule and the soluble GFP-R11 molecules. When the magnetic tweezer exerted around 20 pN of force on the beads, a strong GFP signal was detected colocalising with the SYPRO Orange tethered molecule (Fig.5F). This coalescence was more clearly observed using HILO, likely because the complex is forming >150 nm above the surface distance and is approaching the limit of the TIRF field. The use of fluorescence techniques with magnetic tweezers clearly demonstrates that each tethered R9-R12, once mechanically unfolded, recruits many copies of the GFP-R11 from the solution.

### Vinculin is a mechano-chaperone

Our work has defined that Helix50 is a multi-functional region of talin. When R11 unfolds, this helix is a high affinity vinculin-binding site, but when stretched to an unfolded polypeptide, the VVLINAV region creates a high affinity APR that mediates talin self-assembly. This indicated that the conformation of H50 dictates the interactions it forms, as a helix it binds to vinculin, and as a sheet it binds to other copies of talin. As the APR region in H50 overlaps with the VBS it indicated that these two interactions are mutually exclusive (Fig.6A). Furthermore, it suggested that vinculin might outcompete talin-talin interactions and in doing so resolve these talin aggregates.

**Figure 6.**
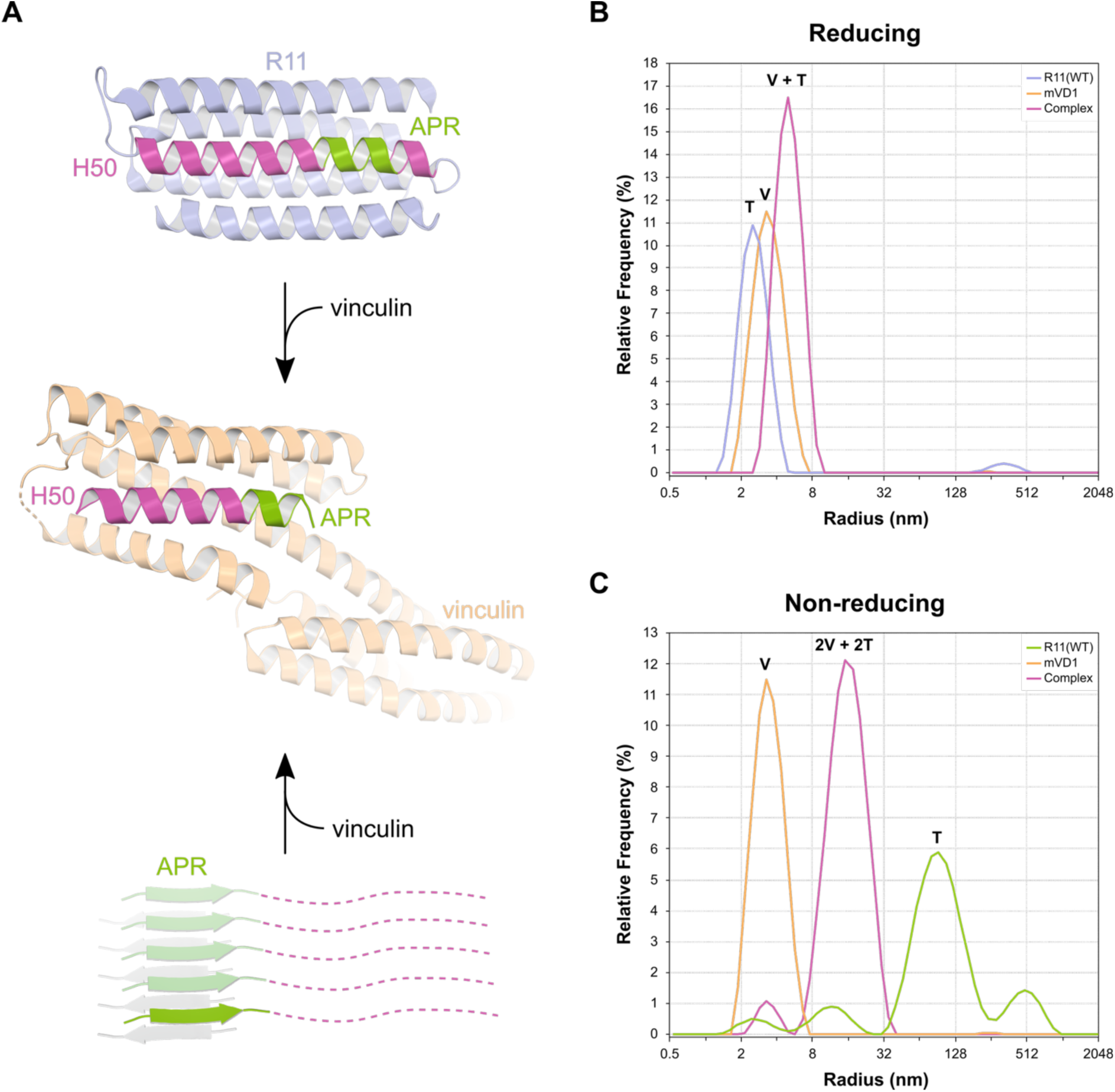
Vinculin acts as a mechano-chaperone for talin. **(A)** R11 (blue) contains a VBS (H50; magenta) in which the APR (green) is also located. Vinculin (orange) can bind to the VBS in both folded and oligomerised states to form a high affinity complex. R11 structure from 3dyj (Gingras et al., 2009), showing residues 2074 – 2140. VD1 in complex with H50 from 4dj9 (Yogesha et al., 2012). **(B-C)** DLS data showing complex formation between R11(WT) and VD1. **(B)** In reducing conditions VD1 binds to H50 when R11 is in the unfolded state. Incubation of VD1 (orange) with R11 (blue) causes a complex (magenta) to form at a larger hydrodynamic radius than either VD1 or R11 alone. **(C)** VD1 can disassemble the soluble aggregates. In non-reducing conditions R11 (green) forms large species, which when mixed with VD1 (orange) results in a complex (magenta) with smaller hydrodynamic radius than R11 alone. The complex peak has a larger hydrodynamic radius than in **(B)** as it consists of an R11 dimer bound to two VD1s. See also <u>Supplementary Movie 1</u>.

To test this, we used DLS (Fig.6B,C), first in the presence of TCEP (Fig.6B) where the DLS profiles showed single peaks for talin R11 (blue), vinculin D1 (orange) and the resultant 1:1 complex (magenta) that forms when they are mixed. In contrast, in the absence of reducing agent (Fig.6C) talin R11 alone forms the big species (as seen in Fig.1B), but this big species is completely lost when mixed with vinculin D1 (magenta). We note here the increased size of the R11-VD1 complex relative to the equivalent peak in Fig.6B, which is due to R11 being a disulphide-linked dimer and bound to two VD1 molecules. However, the DLS unequivocally shows that vinculin has a previously unrecognised chaperone ability for talin as it can resolve these large species.

## Discussion

The challenge with studying potential integrin-binding proteins in cells is complicated as targeting to integrin adhesion complexes can either be via a direct interaction, as is the case with proteins like kindlin and talin, or it can be indirect via interaction with kindlin or the talin scaffold. Here we report that the region of talin previously described as integrin binding site 2 (IBS2) is a high-affinity talin self-interaction site, and that its recruitment to integrin adhesion complexes is predominantly mediated via a direct interaction with talin already at the integrin site. Deletion of R11 from talin at FAs abolishes FA-targeting of GFP-R11 constructs (Fig.2), indicating that the interaction is not via the integrins.

We present data that shows functional aggregation of talin, mediated via a cryptic aggregation prone region (APR), buried within the folded R11 bundle. Intriguingly, the APR region in R11 was previously mapped to be the second integrin-binding site (IBS2), and its localisation to integrin adhesion complexes was concluded to be due to direct binding to integrin beta cytoplasmic tails. We propose that the cryptic binding site in R11 be renamed from IBS2 to APR1 to better reflect its function as a talin-binding site.

### A context dependent binary switch

The talin rod domains serve as force-dependent binary switch domains, as tension can convert them from folded “0” state to unfolded “1” state. This process is reversible when tension goes away and each state has different functions. These binary switches form a central part of the integrin adhesion complexes in most cells. Strikingly, when the R11 helical bundle unfolds, it reveals Helix50, which itself has multiple functions. If vinculin is present, Helix50 can mediate a high affinity connection to the actin cytoskeleton. Force on α-Helices can unravel them and convert them to β-sheets (Minin et al., 2017) and in the context of talin, if vinculin is not available, tension on Helix50 will convert it to a β-sheet conformation stabilising the high affinity talin binding site for other talins to bind to. At the single-molecule level we show that a single stretched talin molecule can unfold other R11 domains to form multimers (Fig.5). This is still a binary switch domain, as the ligands R11 can bind in the unfolded state (vinculin and talin) both require the R11 domain to be unfolded. The discovery of a cryptic talin-binding site that is exposed when R11 is in the unfolded “1” state and unbound to vinculin indicates a context dependence to what the switch engages and further expands the complexity and sophistication of mechanical signalling at adhesion sites. This interdependence between force and ligand availability represents another example of an “AND/OR” logic gate in talin and further expansion of a talin code (Goult et al., 2021). Talin is a synaptic scaffold in the brain and coordinates signalling via its binary switch domains (Ball et al., 2024; Barnett and Goult, 2022; Goult, 2021) so understanding the roles of each switch and the logic underpinning their functioning is crucial to understand mechanical signalling and computation in the brain.

### Harnessing protein unfolding and aggregation in mechanotransduction

The role of protein unfolding in mechanotransduction is now firmly established (Goult et al., 2022; Hytönen and Wehrle-Haller, 2016). Integrin, force exerted on talin, and vinculin have all been shown to ‘trigger’ conformational changes in talin (del Rio et al., 2009; Hytönen and Vogel, 2008; Vigouroux et al., 2020; Yao et al., 2016). However, as mechanotransduction involves unfolding of talin rod domains it raises an intriguing question: what happens to the exposed hydrophobic residues normally resident in the domain core? In talin (Yao et al., 2016) and vinculin (Liu et al., 2025) force-dependent domain unfolding/refolding happens with high fidelity which is no mean feat, but with only 13 of the 61 helices in the talin rod having known function (either VBS or AKAP helices) when exposed, this leaves 48 amphipathic helices exposed and potentially unfolded, which indicates that mechanotransduction will lead to a lot of exposed hydrophobic residues.

Often protein unfolding is harmful, as exposure of hydrophobic residues can result in non-specific aggregation, yet somehow the mechanotransduction machinery seems to circumvent this issue. Cells protect themselves from such aggregates, which are well known drivers of disease, by using chaperones that dissolve and refold these aggregates or direct them for destruction (Liberek et al., 2008).

### Vinculin as a mechano-chaperone

As unwanted protein aggregation is undesirable for healthy function, APR sequences are actively protected from aggregation in the cell. This can be by forming part of a protein-interaction surface, or by being buried in the hydrophobic core of proteins. Alternatively, they can be protected by naturally occurring gatekeeper residues (Reumers et al., 2009) that prevent aggregation should these regions become exposed.

In talin these motifs are not protected by gatekeeper residues, but are cryptic, indicating that mechanical exposure of these APR motifs might facilitate functional aggregation. We propose that aggregation of talin, which occurs in response to mechanotransduction-induced unfolding, is harnessed to build adhesion complexes by forming self-interactions, generating an ‘intracellular meshwork’ and in this work we directly link these self-interactions to mechanical forces and the redox environment. Importantly, resolving these aggregates typically requires interaction with specific chaperones (Liberek et al., 2008), and we show that vinculin is able to resolve talin aggregation, identifying a novel ‘mechano-chaperone’ function for vinculin (Fig.6).

### A redox-sensor at focal adhesions?

Lastly, as unfolding of R11 exposes a highly conserved cysteine that stabilises higher order species *in vitro* it indicates a previously unrecognised link of mechanical signalling and redox-signalling.

The old school of thought was that the redox environment of the cytoplasm was homogeneous, but this view has been challenged by development of probes for reactive oxygen species, hydrogen peroxide, etc. Now it is apparent that highly-localised differences in the redox properties of specific cellular environments are present (Kritsiligkou et al., 2023). This diverse redox landscape is driven by local production of reactive oxygen species or absence of reductive systems at these locations. Sophisticated signalling pathways in cells that sense, respond and relay these redox signals are now widely appreciated (Veal and Kritsiligkou, 2024). The use of fluorescent probes that report on hydrogen peroxide/reductive capacity have revealed that the leading edge of migrating cells sees bursts of increased oxidation that alter the dynamics of these processes (Hurd et al., 2012; O’Mara et al., 2025; Pak et al., 2020), and that the cells use these changes to direct their behaviour. With the identification of mechanically-gated cysteine exposure and localized bursts of H_2_O_2_ generation that enable spatially localized redox signalling, it is possible that mechanotransduction is regulated by redox cues. For example, if the conditions are more oxidising then the R11-R11 interactions might be covalently stabilised by transient disulphide bonds and adhesions might be less dynamic with different signalling. We note that 12 of the 13 talin switch domains contain buried cysteines and future work will focus on the redox-regulation of mechanotransduction.

In summary, we present here a new feature of mechanotransduction, mediated by mechanically-gated aggregation prone regions in talin switches. Identification of a gated talin-binding site in talin reveals a new function of talin switches and a potential new link between mechanical- and redox-signalling.

## Methods

### Peptides, plasmids and cloning

Synthetic peptides of talin1 VVLINAV (2078-2084) and gatekeeper mutant, KVLINAV were ordered from GLBiochem (Shanghai).

The R11(V2078K) and R11(C1978A)-Cys constructs were ordered as synthetic genes in pET151d-TOPO (GeneArt; Life technologies). R11(ΔH47) was subcloned into the XmaI/SacI sites of pET47b using ligation-independent cloning (LIC). GFP-R11 was subcloned into the XmaI/SacI sites of a modified pET47b (eGFP(A206K) inserted between the His-Tag and HRV-3C sites) using LIC. R11(H50) gatekeeper constructs (V2078K, V2078K/V2079K and V2078K/N2082K) were ordered as synthetic genes (GeneArt; Life Technologies) and subcloned into a modified eGFP-C1 vector (MCS replaced with a LIC compatible MCS) using LIC.

### Protein expression and purification

R11 and GFP-R11 constructs were expressed in BL21(DE3). Cells were grown in LB supplemented with 100 µg/mL ampicillin at 37°C. Protein expression was induced with 0.5 mM IPTG at OD_600_ ∼ 0.6 and cells were incubated overnight at 20°C. Cells were harvested by centrifugation, resuspended in lysis buffer (50 mM Tris pH 8, 250 mM NaCl, 5% v/v glycerol) and stored at −20°C.

Proteins were purified using Nickel-affinity, followed by anion-exchange chromatography. Briefly, cells were thawed and supplemented with 1 mM PMSF, 1 mM TCEP and 0.2% v/v Triton X-100. Cells were lysed by sonication and cell debris removed by centrifugation. The supernatant was applied to a 5 mL HisTrap FF (Cytiva) using an Akta Start (Cytiva). The column was washed with 15 column volumes (CV) of wash buffer (50 mM Tris pH 8, 600 mM NaCl, 30 mM Imidazole, 5 mM MgCl_2_, 5 mM ATP, 5% v/v glycerol, 0.2% v/v Triton X-100) followed by 15 CV of nickel buffer A (20 mM Tris pH 8, 250 mM NaCl). Bound protein was eluted across a 14 CV linear imidazole gradient (0 to 300 mM) followed by 4 CV at 500 mM. Fractions containing the protein of interest were pooled and either diluted with 4.5 volumes 20 mM Tris pH 8 or dialysed against 20 mM Tris pH 8, 50 mM NaCl overnight and applied to a 5 mL HiTrap Q HP (Cytiva). Bound protein was eluted across a 14 CV linear NaCl gradient (0 to 600 mM) followed by 4 CV at 1 M. Fractions containing the protein of interest were pooled and dialysed against PBS pH 7.4 overnight. Protein was concentrated, flash-frozen in liquid nitrogen and stored at −80°C.

### Atomic Force Microscopy (AFM)

The VVLINAV peptides were dissolved in Milli-Q water and allowed to self-assemble at room temperature. The samples were deposited onto freshly cleaved mica surfaces (Agar scientific, F7013) and incubated for 10 min. Fibrils were imaged using a Multimode 8 AFM with a Nanoscope V (Bruker) controller operating under peak-force tapping in the ScanAsyst mode. Height channel images were collected with 2048 × 2048 pixels each.

### Dynamic Light Scattering (DLS)

Protein samples were prepared at 50 µM in PBS pH 7.4 in either the presence (reducing) or absence (non-reducing) of 0.5 mM TCEP and loaded into nanoDSF grade standard capillaries (PR-C002; NanoTemper). DLS data were collected using a Prometheus Panta (NanoTemper), performing 10 high sensitivity scans of each capillary at 25°C.

### Fluorescence Polarisation (FP)

The talin1 R11 H50 WT (C-EDPETQVVLINAVKDVAKALGDLISATKAAAG), V2078K (C-EDPETQKVLINAVKDVAKALGDLISATKAAAG), V2078K/V2079K (C-EDPETQKKLINAVKDVAKALGDLISATKAAAG), V2078K/N2082K (C-EDPETQKVLIKAVKDVAKALGDLISATKAAAG) and the TNS3(692-712) (LDIDQSIEQLNRLILELDPT-C) peptides were synthesised by GLBiochem (Shanghai) with either an N-terminal (talin) or C-terminal (tensin) non-native cysteine. Maleimide-fluorescein dye (Thermo Fisher Scientific) was coupled to the peptide following the manufacturer’s protocol. Assays were performed in triplicate with 500 nM peptide and a 2-fold serial dilution of protein in PBS pH 7.4, 0.01% v/v Tween-20, 0.5 mM TCEP. Fluorescence polarisation was measured using a Hidex Sense plate reader (Hidex) at 25°C (excitation: 485 ± 10 nm; emission: 535 ± 20 nm). Data were analysed using OriginPro (OriginLab Corporation) and K_D_ values were generated using a single site with non-specific binding equation.

### Cell culture

*Tln1^-/-^Tln2^-/-^* mouse kidney fibroblast (MKF) cells (Theodosiou et al., 2016) were used and maintained in a humified condition at 37°C and 5% CO_2._ High glucose Dulbecco’s modified Eagle medium (DMEM) supplemented with 10% fetal bovine serum (FBS) was used. The cells were routinely tested for mycoplasma contamination.

### Transfection and expression constructs

Cells were transfected with 1 µg/ 2 kb of plasmid DNA per 10^6^ cells by electroporation using Neon NxT transfection system (Thermo Fisher Scientific). The electroporation parameters were 1350 V, 30 ms, one pulse per 10^6^ cells.

#### Talin expression constructs

##### Full-length constructs

Talin 1–2541, C-terminal mCherry / mScarlet

Talin 1–2541 (V2078K), C-terminal mScarlet

##### Truncations

ΔR1–10: 1–490 + 1974–2541, C-terminal mCherry

ΔR1–12: 1–490 + 2296–2541, C-terminal mCherry

##### Domain fragments

R11: 1974–2140, N-terminal mEGFP VVLINAV: 2078–2084, N-terminal mEGFP

Helix-50: 2072–2104, N-terminal mEGFP

Helix-50 mutants: N-terminal EGFP V2078K; V2078K/V2079K; V2078K/N2082K

### Migration and wound closure assay

Transfected cells were incubated overnight to allow recovery post-transfection. Subsequently, cells were trypsinised and seeded onto 12-well plates coated with 15 µg/mL fibronectin and incubated for 90 min to allow the transfected cells to attach. Following incubation, the medium was replaced with fresh DMEM+10% FBS to remove non-transfected and non-adherent cells. Time-lapse imaging was performed for 24 h at 10 min intervals using a Nikon Eclipse Ti2 fluorescence microscope. Cell migration rate was measured by ImageJ (Fiji) using the MTrackJ plugin (Meijering et al., 2012; Schneider et al., 2012). For wound closure assays, transfected cells were seeded on 24-well plates coated with 15 µg/mL fibronectin and incubated for 48 h to reach the maximum cell confluency. Artificial wounds were created using 10 μL pipette tips and washed with PBS to remove dead and detached cells. Wound healing was monitored by time-lapse imaging for 24 h at 10 min intervals using Nikon Eclipse Ti2 Fluorescence Microscope. Distance travelled was measured using ImageJ.

### Immunostaining and imaging

Transfected cells were seeded on 15 μg/mL fibronectin coated, 8 well glass bottom μ-slides (ibidi) and incubated for 24 h. Following incubation, cells were fixed with 4% paraformaldehyde. Fixed cells were then labelled with phosphorylated paxillin (pY31 paxillin), phosphorylated focal adhesion kinase (pY397 FAK) and activated β1-integrin antibodies, followed by fluorescently labelled secondary antibodies according to the standard immunostaining protocol. Cells were imaged with a Total Internal Reflection Fluorescence (TIRF) microscope (Evident Olympus IX83) using a 100x oil objective. Fluorescence signals were obtained from excitation wavelength 488 nm (GFP, Alexa Flour 488) and 561 nm (mCherry, mScarlet). Images were captured from both epifluorescence (EPI) and TIRF configurations. Laser power and exposure time were adjusted to reduce photobleaching and maintain optimal signal to noise ratio.

### Image analysis

For adhesion segmentation we used the WEKA trainable segmentation plugin (Arganda-Carreras et al., 2017) and classifier to detect focal adhesions (Tinevez et al., 2021) in ImageJ. We selected all adhesions above the size of 5 pixels and quantified the fluorescence intensity to obtain the ratio between talin and protein of interest within adhesion sites.

Adhesion distance from the cell edge was determined by first selecting the cell outline in ImageJ. A Euclidean Distance Map (EDM) was then generated within the cell interior using the Distance Map function in ImageJ (Atherton et al., 2022). Adhesion segmentation was performed as described above. The mean intensity of adhesions mapped onto the EDM was used to quantify their relative position within the cell.

### Single-Molecule Manipulation

These experiments were performed as previously described (Yao et al., 2016), using a custom-built high-force magnetic tweezers instrument capable of applying forces up to 100 pN with an extension resolution of approximately 1 nm (Chen et al., 2011; Zhao et al., 2017). Talin R9–R12 fragments were immobilized on glass coverslips in a laminar flow chamber via HaloTag/Halo-ligand chemistry and tethered to 3 µm paramagnetic beads through biotin–streptavidin interactions. To prevent flow-induced unfolding of the R9–R12 construct during solution exchange of R11, a buffer-isolation membrane well array was employed (Le et al., 2015). The experiments were carried out at a fixed loading rate of 2 pN/s. The force-calibration was based on a standard calibration curve (Zhao et al., 2017). In experiments, the curve requires a single reference point of force versus magnet-bead distance data. The experiments used the well-known DNA overstretching transition (Smith et al., 1996) as the reference point. The resulting overall uncertainty of force-calibration is less than 5%.

### Single-molecule detector construction

The recombinant protein constructs were designed to consist of two R11 domain repeats (or the mutant R11ΔH51) connected by a long, flexible linker sequence. This insert was cloned into a modified pET151 plasmid (6His-AviTag-2xI27-Intein-2xI27-SpyTag) using seamless ligation. All constructs were verified by Sanger sequencing.

The final sequence of R11-L-R11 (coloured as in Fig.5C,D) is 6His-Avi-2XI27-R11-Long Linker-R11-2XI27-SpyTag: HHHHHHGKPIPNPLLGLDSTENLYFQGIDPFTGLNDIFEAQKIEWHEGGGSGLIEVEKPLYGVEVFVGETAHFEIELSEPDVHGQWKLKGQPLAASPDAEIIEDGKKHILILHNAQLGMTGEVSFQAANTKSAANLKVKELGGGSGLIEVEKPLYGVEVFVGETAHFEIELSEPDVHGQWKLKGQPLAASPDAEIIEDGKKHILILHNAQLGMTGEVSFQAANTKSAANLKVKELGGGSGKLGNRGTQACITAASAVSGIIADLDTTIMFATAGTLNREGAETFADHREGILKTAKVLVEDTKVLVQNAAGSQEKLAQAAQSSVATITRLADVVKLGAASLGAEDPETQVVLINAVKDVAKALGDLISATKAAAGKVGDDPAVWQLKNSAKVMVTNVTSLLKTVKAVEDEATGGGSGEFPPAPPLPGDSGTIIPPPPAPGDSTTPPPPPPPPPPPPPLPGGVCISSPPSLPGGTAISPPPPLSGDATIPPPPPLPEGVGIPSPSSLPGGTAIPPPPPLPGSARIPPPPPPLPGSAGIPPPPPPLPGEAGMPPPPPPLPGGPGIPPPPPFPGG PGIPPPPPGMGMPPPPPFGFGVPAAPVLPGSGGGSGGNRGTQACITAASAVSGIIADLDTTIMFATAGTLNREGAETFADHREGILKTAKVLVEDTKVLVQNAAGSQEKLAQAAQSSVATITRLADVVKLGAASLGAEDPETQVVLINAVKDVAKALGDLISATKAAAGKVGDDPAVWQLKNSAKVMVTNVTSLLKTVKAVEDEATLEGGGSGLIEVEKPLYGVEVFVGETAHFEIELSEPDVHGQWKLKGQPLAASPDAEIIEDGKKHILILHNAQLGMTGEVSFQAANTKSAANLKVKELGGGSGLIEVEKPLYGVEVFVGETAHFEIELSEPDVHGQWKLKGQPLAASPDAEIIEDGKKHILILHNAQLGMTGEVSFQAANTKSAANLKVKELGGGSGAHIVMVDAYKPTK

The H51 sequence is underlined and this is deleted in the R11^Δ50^-L-R11^Δ50^ construct. The long flexible linker (182 aa) is from FH1 and a GGGSG short linker was inserted between each component. Some additional residues at the seams are the enzyme digestion sites.

The confirmed plasmids were co-transformed with a secondary plasmid encoding the biotin ligase BirA into *E.coli* BL21(DE3) competent cells. Protein expression was induced in the presence of D-biotin to enable site-specific biotinylation of the AviTag by BirA. Following cell lysis, the 6xHis-tagged fusion proteins were purified using immobilized metal affinity chromatography. The final protein constructs were buffer-exchanged and stored in −80°C for single-molecule experiments.

### TIRF-Magnetic tweezer single molecule fluorescence experiment

Imaging was performed on the iLAS2 Total internal reflection fluorescence (TIRF) system, based on a Nikon Eclipse Ti-2 inverted microscope (Nikon Instruments) equipped with a motorized TIRF illuminator (GATACA systems), a scientific CMOS camera (ORCA-Fusion BT, Hamamatsu), controlled by MetaMorph software (Molecular Devices) and Modular V2.0 GATACA software. Fluorescence images were acquired at 0° (normal) under widefield illumination and at 56-58° incidence angle under Highly inclined and Laminated Optical sheet (HILO) illumination, with either 488nm laser (for GFP) or 561nm laser (for SYTOX orange), using a 100x Apo TIRF objective lens with NA1.49.

We integrated the iLAS2_TIRF system with a pre-calibrated magnet, controlled by an MP-285 step motor to apply force to M270 magnetic beads. The channel treatment process and the working buffer remained consistent with the protocol detailed in the Single-molecule manipulation section.

Moreover, we introduced a 3-kbp DNA handle to act as a spacer between the magnetic bead and the surface. The synthesis of the DNA handle was carried out as described previously (Sun et al., 2024). Subsequently, the DNA handle was covalently attached to the M270 beads (Dynabeads M270-epoxy) via a thiol-epoxy reaction with the epoxy surface. The opposite end of the DNA handle was linked to the Talin R9-R12 AVI-tag via streptavidin.

In preparation for the experiment, 10 μM GFP-H50 was pre-incubated with talin R9-R12 overnight at room temperature. To facilitate the visualisation of the talin R9-R12 location under the bead, 50 nM SYTOX Orange DNA dye (Thermal, S11368) was introduced during the experiment to label the DNA handle.

The imaging processing was performed using ImageJ software.

## Acknowledgements

B.T.G. and N.J.B. were supported by Cancer Research UK Program Grant (CRUK-A21671) and British Heart Foundation Special Project Grant (SP/F/23/150045). B.T.G. and K.B.B. were supported by BBSRC grants BB/S007245/1 (and paired grant BB/S007318/1 to N.H.B.) and BB/N007336/1. V.P.H. acknowledges Research Council of Finland (363941), Sigrid Juselius Foundation and Cancer Foundation Finland for financial support. We acknowledge Biocenter Finland and Tampere Imaging Facility (TIF) for infrastructure support.

We acknowledge Yasumi Otani and Till Kallem for technical assistance and critical reading of the manuscript, and Wei-Feng Xue for assistance with the Atomic Force Microscopy. We thank Pari Kritsiligkou for helpful comments.

## Supplementary Figures

**Supplementary Figure 1.**
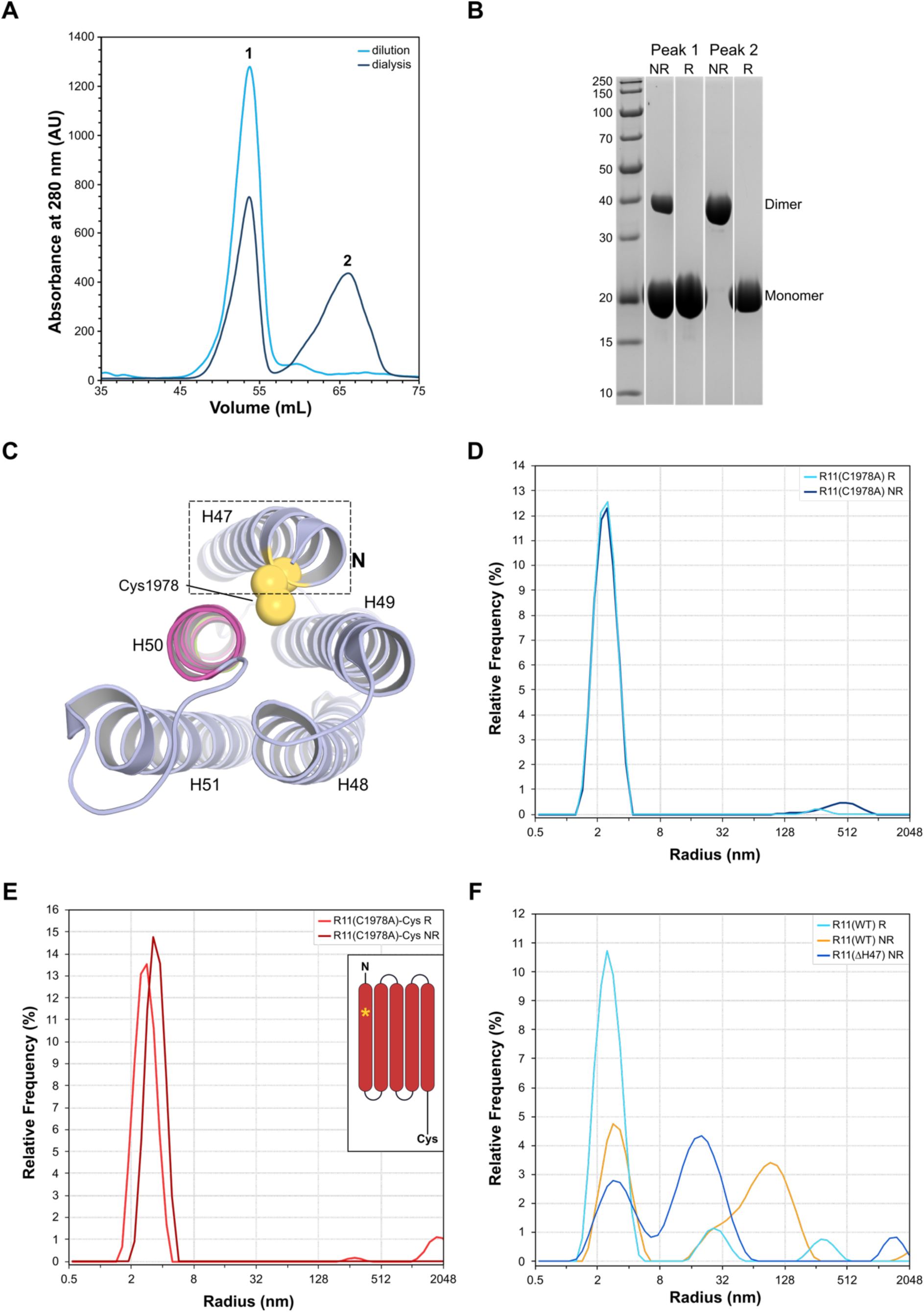
Talin R11 can form big species in solution. **(A)** Two anion exchange chromatography traces of R11. When dialysed overnight, the R11 eluted as two distinct peaks (dark blue). Omitting the dialysis stage resulted in a single peak (cyan). **(B)** Analysis of the anion exchange peaks by non-reducing (NR) and reducing (R) SDS-PAGE highlights that peak 1 contains a mixture of monomeric and disulphide crosslinked R11, whereas peak 2 is solely composed of disulphide-linked R11. **(C)** Talin R11 contains a single buried cysteine residue, Cys1978 in the first helix (H47). **(D-E)** The C1978A mutant of R11 does not form large species and dimerisation itself is not sufficient. DLS intensity distribution of **(D)** R11(C1978A) and **(E)** R11(C1978A)-Cys. The R11(C1978A) mutation does not form higher-order species in either reducing (light blue) or non-reducing (dark blue) conditions. In contrast, R11(C1978A) with a cysteine placed after a short linker at the C-terminus of R11 (schematic inset) can dimerise but does not form higher-order species in either reducing (light red) or non-reducing (dark red) conditions. This demonstrates that the higher-order species seen with R11(WT) is significantly larger than a dimer and that the formation of a disulphide-linked dimer alone is insufficient for formation of a high-order species. **(F)** R11(ΔH47), which lacks the first helix containing the cysteine, forms both monomer and higher-order species even in non-reducing buffer.

**Supplementary Figure 2.**
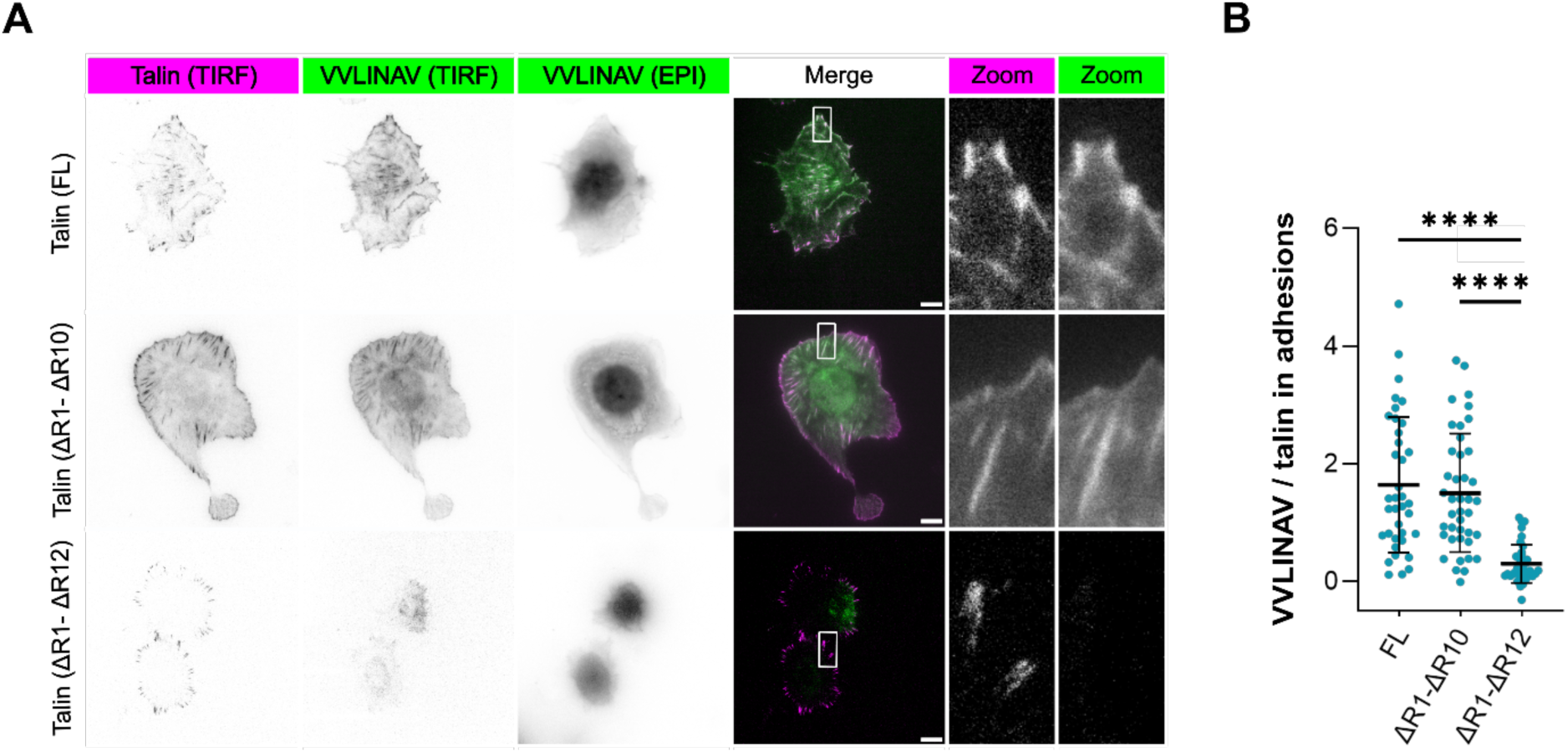
The VVLINAV region of R11 is sufficient to recruit talin to adhesions that contain R11. **(A)** Representative TIRF images showing colocalisation of the VVLINAV region with full-length talin and talin lacking domains R1–R10, Talin(ι1R1-R10) but not with talin lacking domains R1-R12, Talin(ΔR1–R12). Scale bar 10 µm. **(B**) Quantification of VVLINAV colocalisation at adhesion sites for the indicated talin constructs (two independent experiments n ≈ 40 cells). Statistical significance was assessed using Brown–Forsythe and Welch ANOVA tests. Data are presented as mean and SD from three independent biological replicates. Statistical significance is indicated as follows: *P < 0.05; **P < 0.01; ****P < 0.001.

**Supplementary Figure 3.**
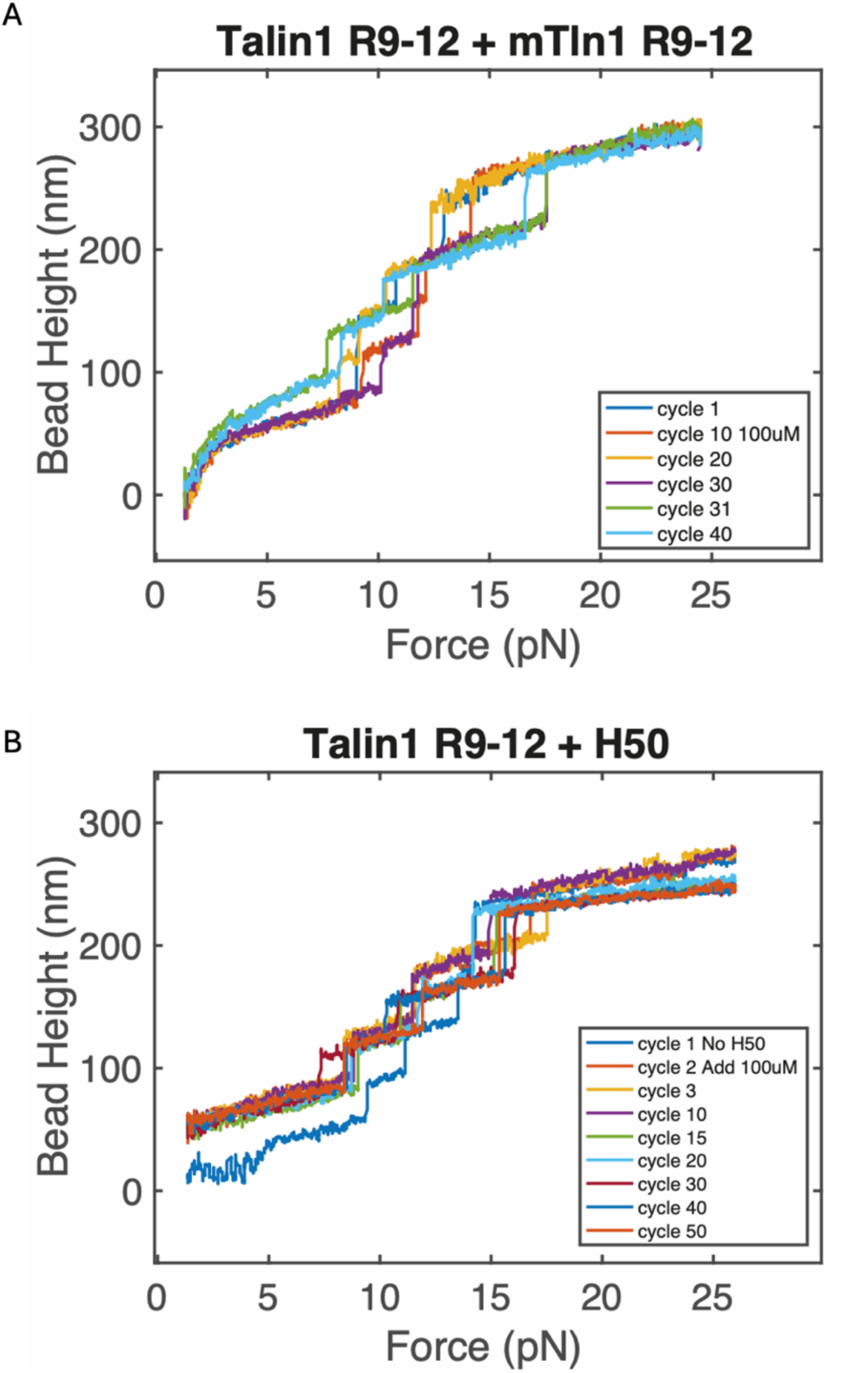
R9-R12 and H50 both bind to mechanically-stretched R9-R12. Representative consecutive force-height curves obtained from a tether of R9-R12 at a loading rate of 0.4 ± 0.04 pN/s are shown. **(A)** 100 µM R9-R12 is added as a ligand to the tethered R9-R12 after cycle 10. Within a few cycles one unfolding step is lost, once lost this step is lost in all subsequent cycles. **(B)** 100 µM H50 is added as a ligand to the tethered R9-R12, and within a few cycles one unfolding step is lost, once lost this step is lost in all subsequent cycles.

## Supplementary Movie

<u>Supplementary Movie 1</u>. Animated version of Figure 6A showing how **Vinculin acts as a mechano-chaperone for talin.** When R11 is folded, H50 and the APR are cryptic inside R11 which is able to bind to “1” state binders like tensin-3. Force on talin leads to unfolding of R11 exposes the H50 which can either bind to vinculin, or be extended to a beta-sheet form where it presents the high affinity Aggregation Prone Region 1 (APR1) which can bind to other talin molecules. Vinculin can resolve these talin multimers by binding to the VBS in H50 demonstrating a novel role of vinculin as a mechano-chaperone for talin.

